## Supplementary Tables and Figures for "Critical factors for precise and efficient RNA cleavage by RNase Y in *Staphylococcus aureus*"

Table S1, list of strains.
Table S2, list of oligos.

#### Supplementary Figures

Figure S1, Alignment of *S. aureus* and *B. subtilis* RNase Y.
Figure S2, PNPase degrades the upstream cleavage fragment of the pSaGap transcript.
Figure S3, EMOTE data from the ΔY strain corresponding to Figures 1, 4, 6 and 8.
Figure S4, Translation does not influence RNase Y cleavage.
Figure S5, Mutations upstream of the RNase Y cleavage sites.
Figure S6, Putative hairpin in pBsGln and inversion of the putative hairpin loop in sector IV of pSaGap.
Figure S7, EMOTE data for hairpins with mutated G-C base-pairs.
Figure S8, Extending the hairpin stem does not alter the cleavage position.
Figure S9, mFold predictions of secondary structures surrounding the RNase Y cleavage sites.

### Supplementary Tables

#### Table S1, list of strains:

| Strain name | Description | Parent strain | Reference |
|---|---|---|---|
| *Staphylococcus aureus* strains | | | |
| PR01 (WT) | Derivative of clinical strain SA564, Δ*pyrFE* and restriction deficient | SA564 | (Redder and Linder 2012) |
| PR01-02 (ΔY) | PR01 with a Δ*rny* deletion (*rny* is also known as *cvfA*) | PR01 | (Redder and Linder 2012) |
| L2ALS01 (BsY) | *S. aureus* RNase Y ORF (*rny* gene) replaced with *B. subtilis* RNase Y ORF (C-terminally tagged with Streptavidine-Flag- His6) | PR01 | This work |
| L2ALS11 | pBsCgg | L2ALS01 | This work |
| L2ALS12 | pBsCgg | PR01 | This work |
| L2ALS13 | pBsCgg | PR01-02 | This work |
| L2ALS22 | pSaGap[ΔIII] | PR01 | This work |
| L2ALS23 | pSaGap[ΔIII] | PR01-02 | This work |
| L2ALS32 | pSaGap[ΔI] | PR01 | This work |
| L2ALS33 | pSaGap[ΔI] | PR01-02 | This work |
| L2ALS37 | pSaGap[ΔVI] | PR01 | This work |
| L2ALS38 | pSaGap[ΔVI] | PR01-02 | This work |
| L2ALS42 | pSaGap[ΔVΔVI] | PR01 | This work |
| L2ALS43 | pSaGap[ΔVΔVI] | PR01-02 | This work |
| L2ALS45 | pSaGap | PR01 | This work |
| L2ALS46 | pSaGap | PR01-02 | This work |
| L2ALS54 | pSaGap[ΔIV] | PR01 | This work |
| L2ALS59 | pSaGap[ΔIV] | PR01-02 | This work |
| L2ALS74 | pSaGap[InvStem] | PR01 | This work |
| L2ALS77 | pSaGap[InvStem] | PR01-02 | This work |
| L2ALS80 | pSaGap[ΔIΔVI] | PR01 | This work |
| L2ALS83 | pSaGap[ΔIΔVI] | PR01-02 | This work |
| L2ALS101 | pSaGap[Δ128] | PR01 | This work |
| L2ALS103 | pSaGap[Δ128] | PR01-02 | This work |
| L2ALS105 | pSaGap[ΔIΔII] | PR01 | This work |
| L2ALS107 | pSaGap[ΔIΔII] | PR01-02 | This work |
| L2ALS109 | pSaGap[ΔIΔII+G] | PR01 | This work |
| L2ALS111 | pSaGap[ΔIΔII+G] | PR01-02 | This work |
| L2ALS113 | pBsCggshort | PR01 | This work |
| L2ALS115 | pBsCggshort | PR01-02 | This work |
| L2ALS117 | pSaGap[A264U] | PR01 | This work |
| L2ALS119 | pSaGap[A264U] | PR01-02 | This work |
| L2ALS121 | pSaGap[U280A] | PR01 | This work |
| L2ALS123 | pSaGap[U280A] | PR01-02 | This work |
| L2ALS125 | pSaGap[ΔIV::CggHP] | PR01 | This work |
| L2ALS127 | pSaGap[ΔIV::CggHP] | PR01-02 | This work |
| L2ALS129 | pSaGap[G268C] | PR01 | This work |

| | | | |
|---|---|---|---|
| L2ALS131 | pSaGap[G268C] | PR01-02 | This work |
| L2ALS147 | pSaGln | PR01 | This work |
| L2ALS149 | pSaGln | PR01-02 | This work |
| L2ALS153 | pSaGln | L2ALS01 | This work |
| L2ALS155 | pBsGln | PR01 | This work |
| L2ALS157 | pBsGln | PR01-02 | This work |
| L2ALS161 | pBsGln | L2ALS01 | This work |
| L2ALS163 | pSaGap[G268C,C276G] | PR01 | This work |
| L2ALS165 | pSaGap[G268C,C276G] | PR01-02 | This work |
| L2ALS167 | pSaGap[A264U,U280A] | PR01 | This work |
| L2ALS168 | pSaGap[A264U,U280A] | PR01-02 | This work |
| L2ALS169 | pSaGap[C276G] | PR01 | This work |
| L2ALS170 | pSaGap[C276G] | PR01-02 | This work |
| L2ALS171 | pSaGap[InvStemGC] | PR01 | This work |
| L2ALS172 | pSaGap[InvStemGC] | PR01-02 | This work |
| L2ALS181 | pSaGap[IIIrandom] | PR01 | This work |
| L2ALS212 | pSaGln[GtoC] | PR01 | This work |
| L2ALS214 | pSaGln[GtoC] | PR01-02 | This work |
| L2ALS216 | pSaGln[GtoC,CtoG] | PR01 | This work |
| L2ALS218 | pSaGln[GtoC,CtoG] | PR01-02 | This work |
| L2ALS236 | pBsCgg[GtoC] | PR01 | This work |
| L2ALS238 | pBsCgg[GtoC] | PR01-02 | This work |
| L2ALS242 | pBsCgg[GtoC,CtoG] | PR01 | This work |
| L2ALS244 | pBsCgg[GtoC,CtoG] | PR01-02 | This work |
| L2ALS248 | pSaGap[ΔV] | PR01 | This work |
| L2ALS250 | pSaGap[ΔV] | PR01-02 | This work |
| L2ALS262 | pSaGap[NoStart] | PR01 | This work |
| L2ALS263 | pSaGap[NoStart] | PR01-02 | This work |
| L2ALS264 | pSaGap[ΔII+G] | PR01 | This work |
| L2ALS265 | pSaGap[ΔII+G] | PR01-02 | This work |
| L2ALS266 | p*fliM*::II-V | PR01 | This work |
| L2ALS267 | p*fliM*::II-V | PR01-02 | This work |
| L2ALS268 | p*fliM* | PR01 | This work |
| L2ALS269 | p*fliM* | PR01-02 | This work |
| **Strain name** | **Description** | **Parent strain** | **Reference** |
| *Bacillus subtilis* strains | | | |
| SSB1002 | Wild-type | | |
| CCB441 | *rny::spc* | SSB1002 | |
| CCB1111 | *amyE::pHM2-cvfA rny:spc* | SSB1002 | |
| CCB1112 | *amyE::pHM2-rny rny::spc* | SSB1002 | |
| *E. coli* strains | | | |
| DH5α | | | Lab strain |
| Stellar | | | Takara |

#### Table S2, list of oligos:

| Oligo number or name | Sequence in 5' to 3' direction (relevant restriction sites have been underlined) | Used for |
|---|---|---|
| | **pSaGap construct** | |
| 337 | TACGAGTCGACGATTAGCGAGTCAGTATAAG | pSaGap construct |
| 58 | TACTACGGCGCGCCTTAACCACCATCAACTACCTCT | |
| 30 | ATTATTGGTACCGACCTTGAATCAAAAGACTT | |
| 29 | TTATCTGGTACCGTCTTTCACTACTACCTCCTC | |
| | **pBsCgg construct** | |
| 26 | AAAAAAGGTACCGCGGTAGCGGGCGGATCA | pBsCgg construct |
| 76 | CAACAAGGCGCGCCATCGTGGTTAGCCGCATCG | |
| 53 | AAAAAAGGTACCAATATCGACCAGGTTCTGTTCC | |
| 50 | AAAACTGTCGACTGGCGCTTTCTGTATAAGA | |
| 431 | TAAATATCTCTCACTTATTTAAAGGAGG | |
| 432 | ACTTCTTTGCGGCTCC | |
| 556 | ATTATTCATGGGTTAATAACTTCTTTGCGGCTC | pBsCgg[GtoC] and pBsCgg[GtoC,CtoG] |
| 557 | CCCTCAATATAAATATCTCTCAC | |
| 558 | GGGTCAATATAAATATCTCTCACTTATTTAAAG | |
| | **pSaGln construct** | |
| 559 | GTCGACCGTTTTAGAAGTCGAAATCG | pSaGln construct |
| 560 | TTAAATGCTCATAGTAACGTATTTGCCTTG | |
| 561 | ACGTTACTATGAGCATTTAACAACAGATGAAC | |
| 562 | GGCGCGCCTTAATCTGATTCTTCGATACGTAC | |
| 592 | TGCCTGCAGGTCGACCGTTTTAGAAGTCGAAATCGCA | |
| 593 | TAGAATAGGCGCGCCTTAATCTGATTCTTCGATACGTACGAA | |
| 700 | GGAGCATTTATTGGCAAAGTTTCTCC | pSaGln[GtoC] and pSaGln[GtoC,CtoG] |
| 701 | TGATTTATGGGCATTTATTAAATAAAATTTGGAGGATT | |
| 702 | TGATTTATCCCGATTTATTAAATAAAATTTG | |
| | **pBsGln construct** | |
| 565 | ATGTTTCATTGGATAATATCGAATTTGTCTTGCT | pBsGln construct |
| 566 | TTCGATATTATCCAATGAAACATGATCTGTC | |
| 580 | TTTGTCGACTTTCTCTGGATTTG | |
| 581 | TTTGGCGCGCCTTACTCTTC | |
| 582 | TGCCTGCAGGTCGACTTTCTCTGGATTTGATGTTAAGAATCC | |
| 583 | TAGAATAGGCGCGCCTTACTCTTCGATACGAACGAATCC | |
| | **pSaGap mutations** | |
| 333 | TCGTATAGGCGCGCCTTAACCACC | pSaGap[ΔIII] |
| 334 | TCGTAGGTACCGTCTTTCACTACTACC | pSaGap[ΔI], pSaGap[ΔIΔII], pSaGap[ΔIΔII+G] |
| 337 | TACGAGTCGACGATTAGCGAGTCAGTATAAG | pSaGap[ΔIII] |
| 338 | TGAATAAGTAAAAAGTTTAATACTTTTTAAATATC | pSaGap[ΔIII] |
| 339 | AAAGTATTAAACTTTTTACTTATTCAAGTATTATCTTTGCTG | |
| 342 | TACGAGGTACCATACTTGAATAAGAGATAAAAAG | pSaGap[ΔI] |

| | | |
|---|---|---|
| 343 | AATCAGAATTCGTTAAGGCGCGCCTATTCTAAATGC | pSaGap[ΔVI],<br>pSaGap[ΔVΔI] |
| 344 | TACTGAATTCTACCAAAACCATTAATTGCT | pSaGap[ΔVI] |
| 345 | TACTGGAATTCTTAAAAAGTATTAAACTTTTTATCTCTTA | pSaGap[ΔVΔI] |
| 421 | AATATCATTTTAAAGGAGGCCATTATA | pSaGap[ΔIV] |
| 424 | TCTCTTATTCAAGTATTATCTTTGCTG | |
| 423 | AATTGAAAAATAATATCATTTTAAAGGAGGCC | pSaGap[InvStem] |
| 424 | AATGAAAAATTCTCTTATTCAAGTATTATCTTTGCTG | |
| 478 | TACGAGGTACCAATACTTGAATAAGAGATAAAAAG | pSaGap[Δ128] |
| 29 | TTATCTGGTACCGTCTTTCACTACTACCTCCTC | |
| 479 | TTTGGTACCAGATAAAAAGTTTAATACTTTTTAAATATCAT | pSaGap[ΔIΔII] |
| 480 | TTTGGTACCGAGATAAAAAGTTTAATACTTTTTAAATATC | pSaGap[ΔIΔII+G] |
| 513 | ATATCTCTTATTCAAGTATTATCTTTGC | pSaGap[A264U] |
| 514 | AAAGTTTAATACTTTTTAAATATCATTTTAAAG | |
| 515 | TAAAGTATTAAACTTTTTATCTCTTATTC | pSaGap[U280A] |
| 516 | TAAATATCATTTTAAAGGAGGCC | |
| 517 | TAATCCCTCAATATATATCATTTTAAAGGAGGCC | pSaGap[cggRSL] |
| 518 | TTCATCCCTTAATAACTCTTATTCAAGTATTATCTTTGC | |
| 553 | GTTTTTATCTCTTATTCAAGTATTATCT | pSaGap[G268C] |
| 554 | TTTAATACTTTTTAAATATCATTTTAAAGG | |
| 563 | TAAAGTATTAAACTTTATATCTCTTATTCAAG | pSaGap[A264U,U280A<br>] |
| 634 | NNNTAAAAAGTTTAATACTTTTTAAATATCATTTT | pSaGap[IIINNN]<br>(variant) |
| 636 | CTTATTCAAGTATTATCTTTGCTG | |
| 640 | TAAAGTATTAAACTTTATATCTCTTATTCAAG | pSaGap[A264U,U280A<br>] |
| 641 | AATTCAAAAATAATATCATTTTAAAGGAGGC | pSaGap[InvStemGC] |
| 642 | AATCAAAAATTCTCTTATTCAAGTATTATCTTT | |
| 643 | TTTAATAGTTTTTAAATATCATTTTAAAGGAGGC | pSaGap[C276G] |
| 644 | CTTTTTATCTCTTATTCAAGTATTATCTT | |
| 706 | TATCTTTGCTGCGGCTTC | pSaGap[ΔII+G] |
| 707 | GAGATAAAAAGTTTAATACTTTTTAAATATC | |
| 708 | ATATTTAAAAAGTATTAAACTTTTTATCTCTTATT | pSaGap[ΔV] |
| 709 | GGTCGTTTAGCATTCAGAAG | |
| 703 | TAAAAAGACGGTACCGACCTTG | pSaGap[NoStart] |
| 704 | TACTACCTCCTCCTTATATTTATAAATGTAAATAA | |
| 799 | TAATTACTTTTTATCTCTTATTCAAGTATTATC | pSaGap[InvLoop] |
| 800 | ACTTTTTAAATATCATTTTAAAGGAGG | |
| 925 | GCAGCAAAGATAGAGATAAAAAGTTTAATACTT | SecIImutant1 |
| 926 | CTCTATCTTTGCTGCTATCTTTGCTGCGGCTTC | |
| 927 | TGGAGAAACTTTGAGATAAAAAGTTTAATACTT | SecIImutant2 |
| 928 | CTCAAAGTTTCTCCATATCTTTGCTGCGGCTTC | |
| 935 | CTCGCCGAATAAGAGATAAAAAGTTTAATACTT | SecIImutant3 |
| 936 | CTCTTATTCGGCGAGTATCTTTGCTGCGGCTTC | |
| 937 | ATACTTCTCGCCGAGATAAAAAGTTTAATACTT | SecIImutant4 |
| 938 | CTCGGCGAGAAGTATTATCTTTGCTGCGGCTTC | |

| | | |
|---|---|---|
| | **Cloning *fliM gap*** | |
| 733 | GGCTGTTATACTTGAATAAGAGATAAAAAGTTT | *fliM gap* transcriptionnal fusion construct |
| 734 | TCAAGTATAACAGCCCCATACGAAAAT | |
| 735 | TAGAATTCAACCTGCTGCGTCGTAGCC | |
| 736 | GCAGGTTGAATTCTACCAAAACCATTAAT | |
| 737 | GGCGCGCCTATTCTAAATG | |
| 738 | TAGAATAGGCGCGCCAGCGTCCATCGCCGCCA | |
| 739 | GTTAAAGTTTTACCAAGTATTCTTTCTCAAGCTG | |
| 740 | TGGTAAAACTTTAACACAAGCATTAC | |
| 794 | GATAAAAAGTTTAATACTTTTTAAATATCATTT | pSaGap[IIIGGA] |
| 793 | CCTTATTCAAGTATTATCTTTGCTGC | |
| 796 | TAAAAAGTTTAATACTTTTTAAATATCATTTTAAA | Forward for pSaGap[IIIAUU], pSaGap[IIICUA] and pSaGap[IIICGA] |
| 795 | AATCTTATTCAAGTATTATCTTTGCTGC | pSaGap[IIIAUU] |
| 797 | TAGCTTATTCAAGTATTATCTTTGCTG | pSaGap[IIICUA] |
| 798 | TCGCTTATTCAAGTATTATCTTTGCTG | pSaGap[IIICGA] |
| | **Northern blot** | |
| 310 | TTAACTTCTGTGTTCGGCATGGGAACAGGTGTGACCTCC | Northern probe 5S |
| 1 | TCAAAATTATACATGTCAACGA | Northern probe P1 |
| 408 | GTGGTTGCCTTTTTAAGTCCCGCGTGGGAC | Northern probe P2 |
| CC058 (16S) | CAGCGTTCGTCCTGAGCCAG | Northern probe B. subtilis 16S |
| CC2437 (atpB) | CATAGTAGTGGGTTAAAGCAACAAC | Northern probe B. subtilis atpB |
| CC2438 (glnA) | TACACCGAACAATGGTTTTGGCATAA | Northern probe B. subtilis glnA |
| | **EMOTE** | |
| 298 | GGCATTCCTGCTGAACCGCTCTTCCGATCTTACATGTCAACGATAATACA | Reverse transcription in targeted EMOTE (This work) |
| BioRp8 | Biotin-dG-CGGCACCAACCGAGGVVVVVVVACAGA (RNA oligo) | EMOTE Ligation (Redder, 2018) |
| D6A | CTCTTTCCCTACACGACGCTCTTCCGATCTNTACACGGCACCAACCGAGG | Second strand PCR (Khemici et al., 2015) |
| D6B | CTCTTTCCCTACACGACGCTCTTCCGATCTNGTATCGGCACCAACCGAGG | |
| D6C | CTCTTTCCCTACACGACGCTCTTCCGATCTNCGTCCGGCACCAACCGAGG | |
| D6D | CTCTTTCCCTACACGACGCTCTTCCGATCTNAAGTCGGCACCAACCGAGG | |
| D6E | CTCTTTCCCTACACGACGCTCTTCCGATCTNACACCGGCACCAACCGAGG | |
| D6F | CTCTTTCCCTACACGACGCTCTTCCGATCTNGGTACGGCACCAACCGAGG | |
| D6H | CTCTTTCCCTACACGACGCTCTTCCGATCTNTCGGCGGCACCAACCGAGG | |
| D6I | CTCTTTCCCTACACGACGCTCTTCCGATCTNCAAGCGGCACCAACCGAGG | |
| D6J | CTCTTTCCCTACACGACGCTCTTCCGATCTNTTGACGGCACCAACCGAGG | |

| | | |
|---|---|---|
| D6K | CTCTTTCCCTACACGACGCTCTTCCGATCTNGCTGCGGCACCAACCGAGG | |
| D6L | CTCTTTCCCTACACGACGCTCTTCCGATCTNCCGACGGCACCAACCGAGG | |
| D6M | CTCTTTCCCTACACGACGCTCTTCCGATCTNCTCGCGGCACCAACCGAGG | |
| D6N | CTCTTTCCCTACACGACGCTCTTCCGATCTNAGGACGGCACCAACCGAGG | |
| D6O | CTCTTTCCCTACACGACGCTCTTCCGATCTNATTGCGGCACCAACCGAGG | |
| D6P | CTCTTTCCCTACACGACGCTCTTCCGATCTNGACGCGGCACCAACCGAGG | |
| D6Q | CTCTTTCCCTACACGACGCTCTTCCGATCTNTGTTCGGCACCAACCGAGG | |
| A-PE-PCR10 | AATGATACGGCGACCACCGAGATCTACACTCTTTCCCTACACGACG | |
| B-PE-PCR20 | CAAGCAGAAGACGGCATACGAGATCGGTCTCGGCATTCCTGCTGAACCGC | |
| | **DMS-MaPseq** | |
| 835 | gcatctaatacgactcactatggcGTAAAGAGGTTAATTTTTGTCCCACGCGG | PCR to add T7 promoter to 540bp of gap operon. |
| 463 | cagtatttattatgcatttagaataggcgc | |
| 868 | CTGGAGTTCAGACGTGTGCTCTTCCGATCTATGCTAAACGACCAATTCTA | RT-primer for DMS-MaPseq |
| 815 | CTCTTTCCCTACACGACGCTCTTCCGATCTNACAGTCACTGATGAAGCCGCAGC | Second strand synthesis, adding barcode "ACAG" |
| 816 | CTCTTTCCCTACACGACGCTCTTCCGATCTNTACCTCACTGATGAAGCCGCAGC | Second strand synthesis, adding barcode "TACC" |
| 817 | CTCTTTCCCTACACGACGCTCTTCCGATCTNCGTTTCACTGATGAAGCCGCAGC | Second strand synthesis, adding barcode "CGTT" |
| 818 | CTCTTTCCCTACACGACGCTCTTCCGATCTNTCGTTCACTGATGAAGCCGCAGC | Second strand synthesis, adding barcode "TCGT" |
| 233 | AATGATACGGCGACCACCGAGATCTACACTCTTTCCCTACACGACG | PCR amplification, forward primer |
| 255 | CAAGCAGAAGACGGCATACGAGATATTGGCGTGACTGGAGTTCAGACGTGTGC | PCR amplification, reverse primers |
| 259 | CAAGCAGAAGACGGCATACGAGATTGGTCAGTGACTGGAGTTCAGACGTGTGC | |
| 260 | CAAGCAGAAGACGGCATACGAGATCACTGTGTGACTGGAGTTCAGACGTGTGC | |
| | **Sector III cleavage-efficiency quantification** | |
| 762 | CTCTTTCCCTACACGACGCTCTTCCGATCTNTGGCAAAGATAATACTTGAATAAG | Plasmid quantification for the Sector III variant library |
| 763 | CTCTTTCCCTACACGACGCTCTTCCGATCTNGTGCAAAGATAATACTTGAATAAG | |
| 764 | CTCTTTCCCTACACGACGCTCTTCCGATCTNACGCAAAGATAATACTTGAATAAG | |
| 765 | CTCTTTCCCTACACGACGCTCTTCCGATCTNTCGCAAAGATAATACTTGAATAAG | |
| 801 | CTCTTTCCCTACACGACGCTCTTCCGATCTNCAGCAAAGATAATACTTGAATAAG | |
| | **Generating *B. subtilis* mutants** | |
| CC2228 | CATAGCAAGAGGAGGTGAAAGTGAATTTATTAAGCCTCCTAC | Cloning of SaY |
| CC2155 | GATGCTTAGCGCATCACTTTATTTCGCATATTCTACTGCTC | |
| CC2150 | cactaaGAATTCTTGACAAGTATTTCCGAC | Cloning 5'end UTR of BsY |
| CC2229 | GTAGGAGGCTTAATAAATTCACTTTCACCTCCTCTTGCTATG | |
| CC2154 | GAGCAGTAGAATATGCGAAATAAAGTGATGCGCTAAGCATC | Cloning 3'end UTR of BsY |
| CC2151 | CACTAAGTCGACTCTTCTTGAAAATTCCTTG | |

### Supplementary Figures

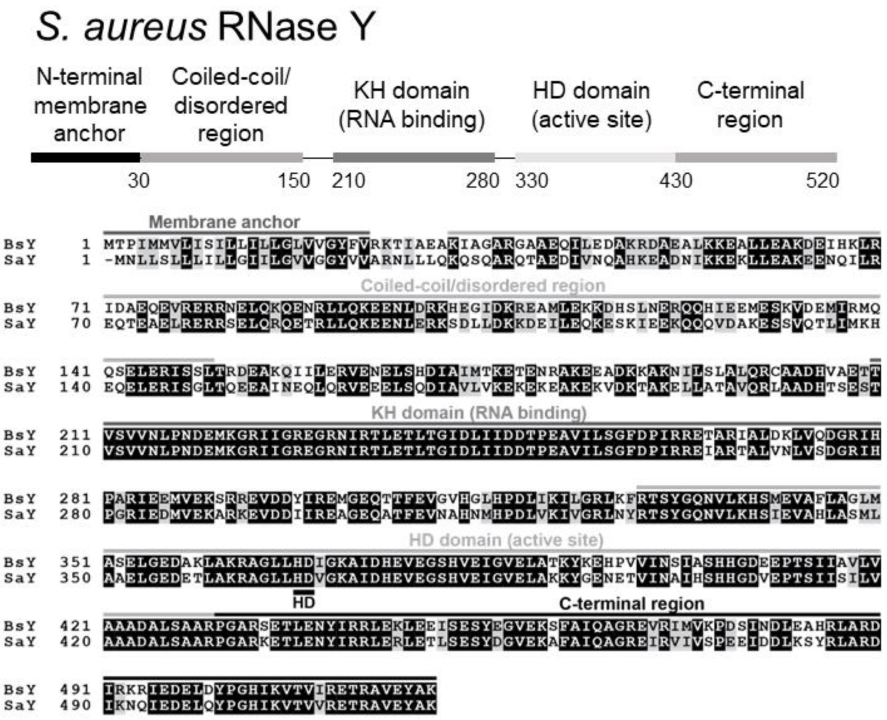


**Figure S1, Alignment of *S. aureus* and *B. subtilis* RNase Y.** SaY: *S. aureus* RNase Y, BsY, *B. subtilis* RNase Y, HD: the two critical amino acids in the active site. Adapted from Redder, 2018.

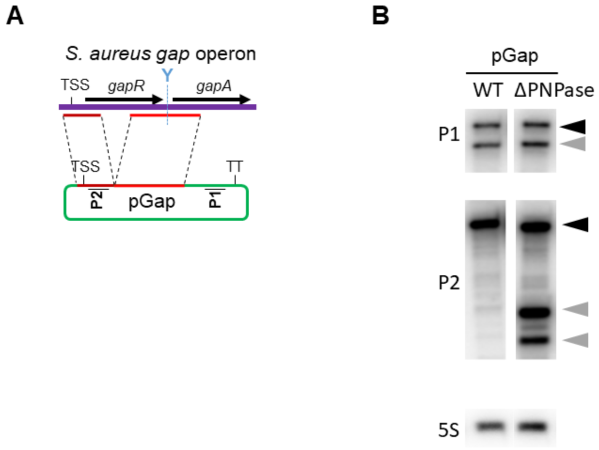


**Figure S2, PNPase degrades the upstream cleavage fragment of the pSaGap transcript.** A) The layout of the pSaGap construct, with the location of the upstream and downstream Northern blot probes (P2 and P1, respectively).

B) Northern blot showing the appearance of two new bands when using the P2 (upstream) probe on total RNA from a PNPase deletion strain carrying pSaGap. Black and grey triangles indicate full-length transcript and cleavage products, respectively.

This data is in agreement with a large-scale *Streptococcus pyogenes* study where upstream products of RNase Y cleavage are systematically degraded by PNPase (Broglia et al. 2020).

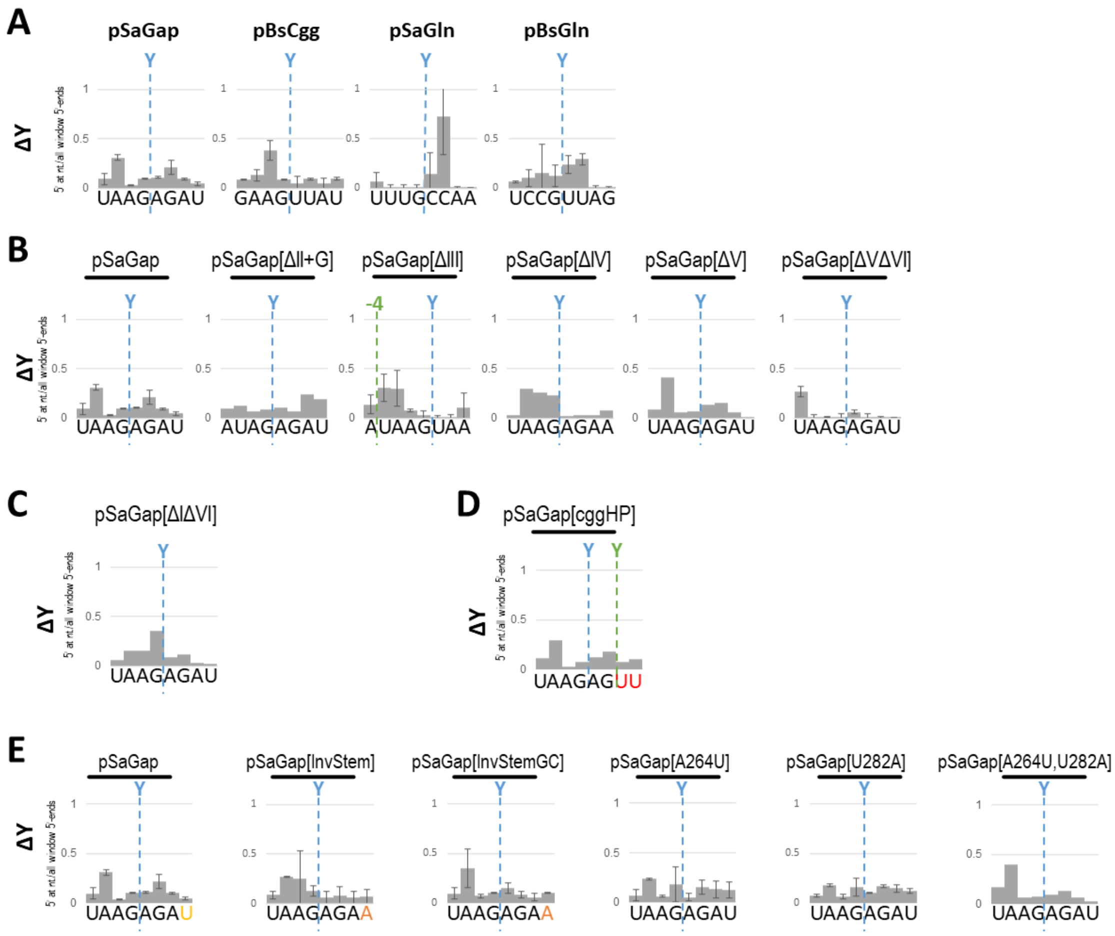


**Figure S3, EMOTE data from the ΔY strain corresponding to Figures 1, 4, 6 and 8.**
The EMOTE data is presented as proportions of RNA molecules with a given 5' end on the Y-axis (number of reads detected at a specific position divided by the total number of reads detected within the chosen window).
Note that the number of detected molecules with 5'ends within the shown windows is much larger in the WT strain than in the ΔY strain, since RNase Y does not cleave the RNAs in the ΔY strain. The ΔY data is therefore based on a very low number of detected molecules and corresponds to background noise. The random nature of this noise can sometimes lead to a tall column for a position, since the EMOTE data is presented as proportions, with sum of the columns set to 1 (this is for example the case for the 6$^{th}$ position in the ΔY pSaGln data on panel A, where no cleavage is observed in the Northern blot in Figure 1C).
The native RNase Y cleavage position is indicated with blue dotted lines. Green dotted lines indicate shifted cleavage positions where relevant. A) Corresponds to Figure 1. B) Corresponds to Figure 4C. C) Corresponds to Figure 4D. D) Corresponds to Figure 6. E) Corresponds to Figure 8.

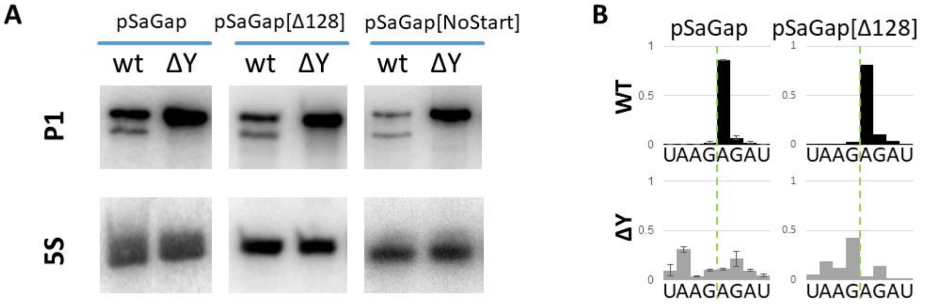


**Figure S4, Translation does not influence RNase Y cleavage.** A) Northern blot showing RNase Y dependent cleavage in pSaGap with a frame-shift (pSaGap[Δ128]) and with the start-codon mutated (pSaGap[NoStart]). B) EMOTE data confirming that the frame-shift in pSaGap[Δ128] has not shifted the RNase Y cleavage position.

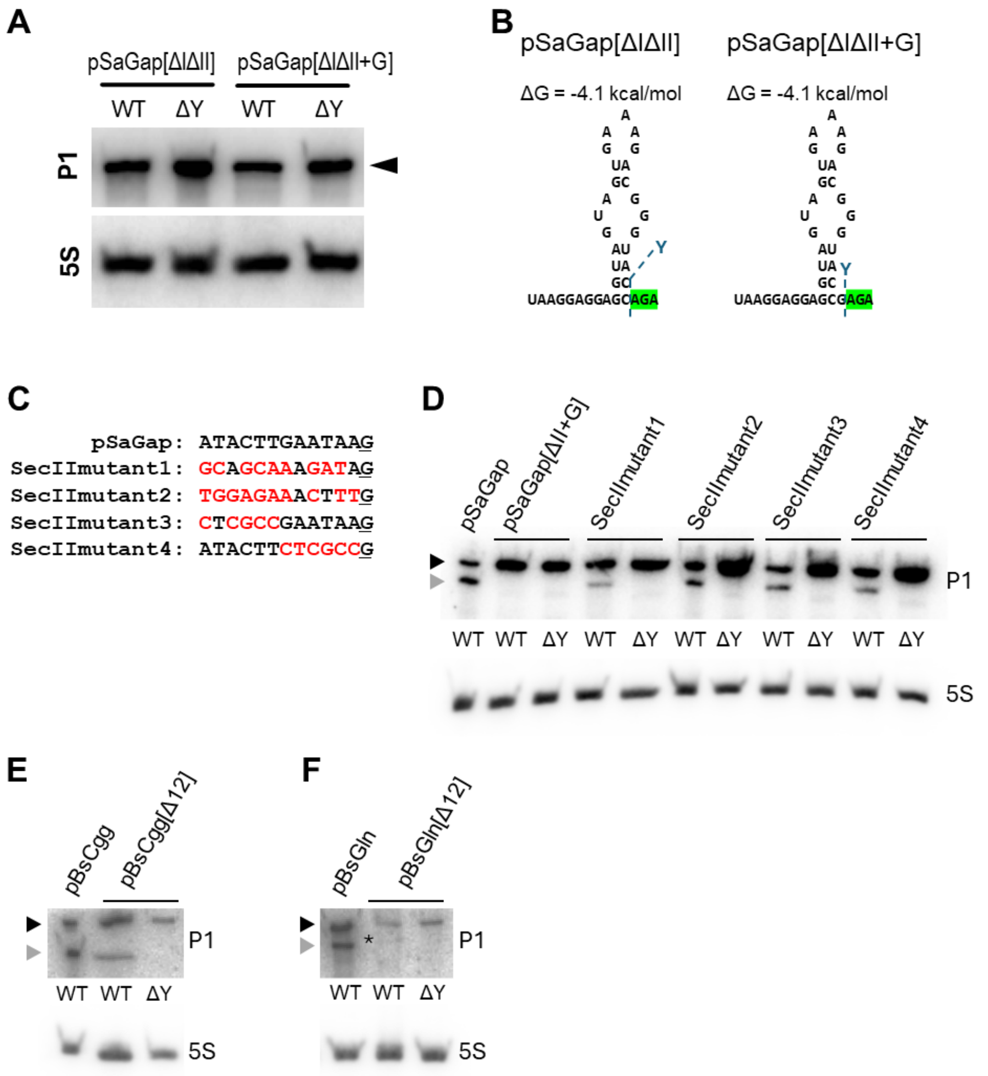


**Figure S5, Mutations upstream of the RNase Y cleavage sites.**
A) Northern blot for the deletions of sectors I and II of pSaGap, with or without the G upstream of the cleavage position.
B) Putative secondary structure that can form if both sectors I and II of pSaGap are deleted. Native RNase Y cleavage positions are indicated with blue dotted lines. Sector III is highlighted in green.
C) The sequences of the wild-type pSaGap sector II and the four mutated versions. The G immediately upstream of the cleavage site has been underlined, and the nucleotides that differ from the wild-type sequence are shown in red.
D) Northern blot showing that while the transcript from pSaGap[ΔII+G] is not cleaved, the transcripts from all four mutant variants are cleaved. The strain background is shown below each lane. The full-length and cleaved transcripts are indicated by the black and grey arrowheads, respectively.
E) Northern blot showing cleavage of pBsCgg and BsCgg[Δ12] (where 12 nucleotides immediate upstream of the conserved G have been deleted).
F) Northern blot showing cleavage of pBsGln and BsGln[Δ12] (where 12 nucleotides immediate upstream of the conserved G have been deleted). A weak band corresponding to the cleaved product has been marked with an asterisk.

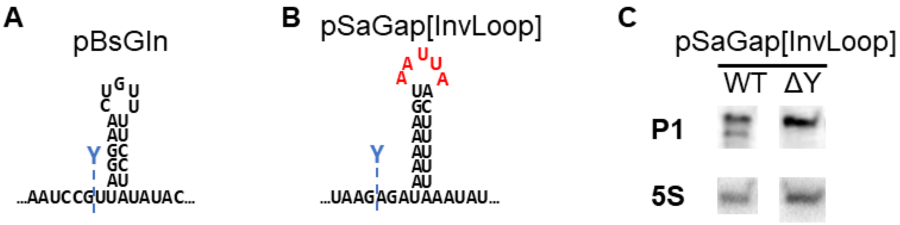


**Figure S6, Putative hairpin in pBsGln and inversion of the putative hairpin loop in Sector IV of pSaGap.**
A) The putative hairpin immediately downstream of the RNase Y cleavage in pBsGln.
B) The sequence of the putative InvLoop hairpin where the loop has been modified so that U becomes A and A becomes U (mutated nucleotides shown in red).
C) Northern blot showing the cleavage of the pSaGap[InvLoop] transcript.

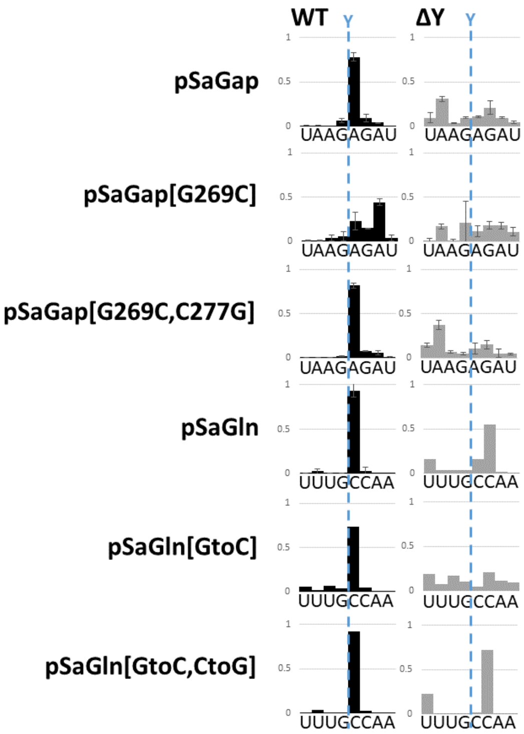


**Figure S7**, **EMOTE data for hairpins with mutated G-C base-pairs, corresponding to Figure 7A and 7C in the main text.** Note that the EMOTE data is presented as proportions of RNA molecules with a given 5' end on the Y-axis (number of reads detected at a specific position divided by the total number of reads detected within the chosen window), and that the number of detected molecules is much larger in the WT strain than in the ΔY strain.

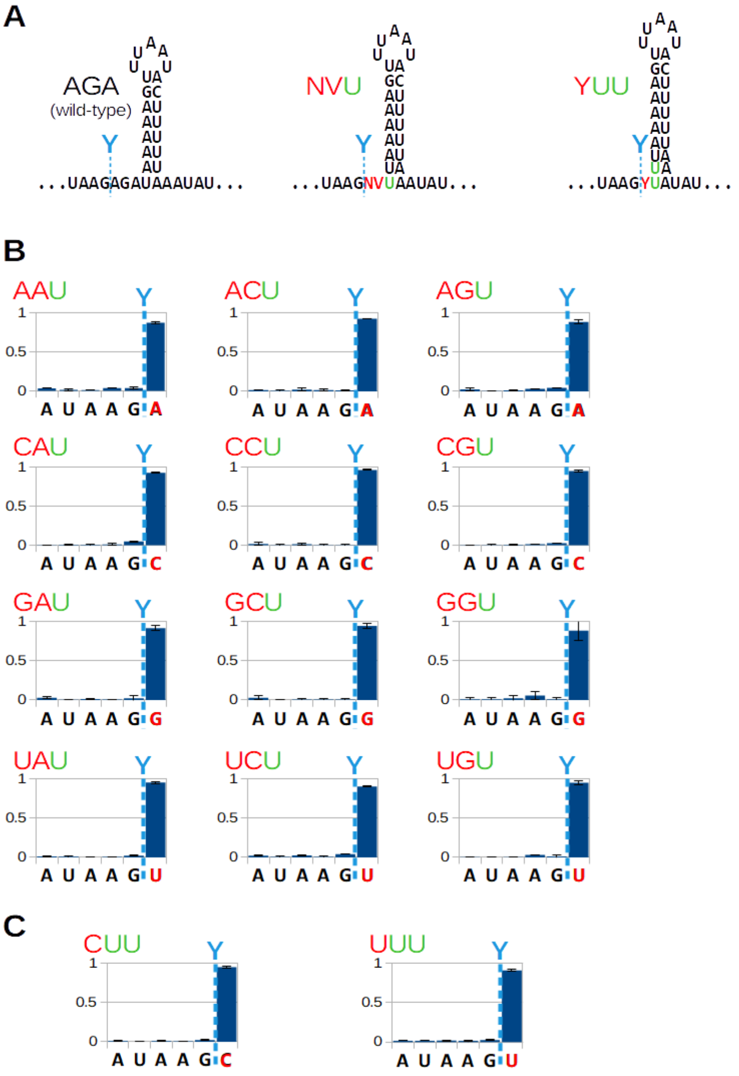


**Figure S8, Extending the hairpin stem does not alter the cleavage position.**
A) Wild-type (AGA) *Sa-gapR* hairpin, hairpin with a single base-pair extension (NVU) and hairpin with two base-pairs extension (YUU). Y: Pyrimidine bases, V: A, C or G. Red nucleotides are varied and green U's are the uridines that extend the putative hairpin stem by one or two base-pairs (NVU and YUU, respectively).
B) EMOTE data from NVU trinucleotide combinations that can potentially extend the stem by a single base-pair. The cleaved nucleotide sequence is shown below the graph and the native cleavage position is shown by a light blue dotted line. Red nucleotides are varied and the green U's extend the putative hairpin stem by one base-pair.
C) EMOTE data from YUU trinucleotide combinations that can potentially extend the stem by two base-pairs. The cleaved nucleotide sequence is shown below the graph and the native cleavage position is shown by a light blue dotted line. Red nucleotides are varied and the green U's are the uridines that extend the putative hairpin stem by two base-pairs.

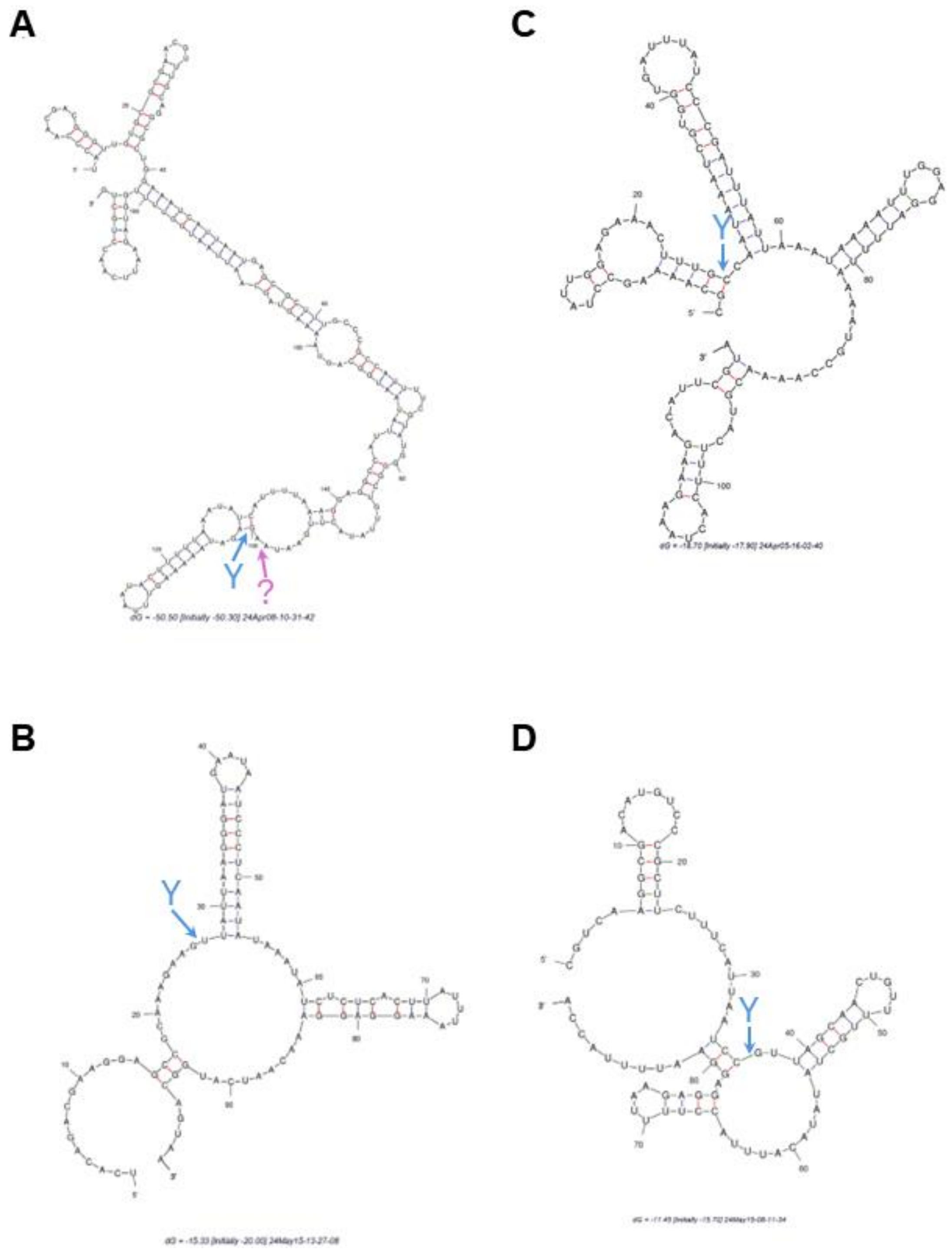


**Figure S9, mFold predictions of secondary structures surrounding the RNase Y cleavage sites.** RNase Y cleavage sites are indicated with blue arrows. Structure prediction and free energy calculations were performed on the mFold.org server using default settings (Zuker 2003).

A) The pfliM::II-V transcript. The RNase Y independent cleavage position is indicated in purple.
B) pBsCgg transcript.
C) pSaGln transcript.
D) pBsGln transcript.

### References for supplementary materials

Broglia L, Lécrivain A-L, Renault TT, et al (2020) An RNA-seq based comparative approach reveals the transcriptome-wide interplay between 3'-to-5' exoRNases and RNase Y. Nat Commun 11:1587. https://doi.org/10.1038/s41467-020-15387-6

Redder P, Linder P (2012) New range of vectors with a stringent 5-fluoroorotic acid-based counterselection system for generating mutants by allelic replacement in Staphylococcus aureus. Appl Environ Microbiol 78:3846–3854. https://doi.org/10.1128/AEM.00202-12

Zuker M (2003) Mfold web server for nucleic acid folding and hybridization prediction. Nucleic Acids Res 31:3406–3415. https://doi.org/10.1093/nar/gkg595
